# Suboptimal foraging decisions and involvement of the ventral tegmental area in human opioid addiction

**DOI:** 10.1101/2022.03.24.485654

**Authors:** Candace M. Raio, Kathryn Biernacki, Ananya Kapoor, Kenneth Wengler, Darla Bonagura, Joany Xue, Sara M. Constantino, Guillermo Horga, Anna B. Konova

## Abstract

Addiction is marked by a tendency to exploit sources of reward despite diminishing returns. This behavior is aptly captured by animal patch-foraging models that have recently been extended to humans. Dopamine and norepinephrine centrally mediate addictive behavior and activity in both catecholaminergic systems is proposed to reflect the computations necessary for optimal foraging. However, the specific neural bases of excessive foraging and their role in human addiction are largely unknown. To address this gap, we studied the behavior of people with and without opioid use disorder (OUD) on a patch-foraging task in which they made serial decisions to “harvest” a depleting resource (“patch”) for reward or incur a varying cost to “travel” to a replenished patch. In a subset of participants, we used high-resolution neuromelanin-sensitive MRI to image neuromelanin concentration, a proxy for long-term catecholaminergic function, in distinct dopaminergic nuclei (ventral tegmental area, substantia nigra subregions) and the noradrenergic locus coeruleus. While all participants were sensitive to the long-run reward rates of different patch-foraging environments, OUD participants stayed in reward patches longer than optimal—markedly overharvesting a source of reward despite its declining value—and this correlated with more chronic drug use. Overharvesting was selectively associated with lower neuromelanin signal in the ventral tegmental area but not other dopaminergic nuclei, nor the locus coeruleus. Our findings suggest that foraging decisions relevant to addiction involve a ventral-tegmental-area circuit that may signal reward rates in dynamic environments and implicate this circuit in maladaptive reward pursuit in human addiction to opioids.

**Significance statement:** Patch-foraging provides a potentially important translational framework for understanding addictive behavior by revealing how maladaptive reward pursuit emerges in more ecologically valid decision contexts. Here, we show that the tendency to exploit sources of reward despite diminishing returns is associated with chronic drug use in people with opioid use disorder, a particularly devastating form of addiction. We further use neuromelanin-sensitive MRI, a neuroimaging measure of the long-term function of dopamine neurons, to reveal that variation in ventral tegmental area neuromelanin signal selectively underlies individual differences in this overharvesting bias. These findings establish a role for specific dopaminergic circuits in patch-foraging decisions and advance understanding of the neurobiology of human addiction to opioids that has so far eluded the field.

## Introduction

A fundamental aspect of maladaptive reward pursuit is the tendency to continue to engage with a particular reward source despite diminishing returns. This is perhaps best exemplified by drug addiction, where individuals persist in drug-seeking behavior even when a drug’s value declines and alternative sources of reward are available (1), and is especially striking in the case of opioid use disorder (OUD) where drug use rates have reached epidemic levels and more people than ever are dying from opioid overdose. Obtaining a better understanding of the behavioral and neural mechanisms that render individuals vulnerable to maladaptive reward behaviors, and to continued drug seeking and use, is thus critical.

Efforts aimed at characterizing this type of reward-related behavior have traditionally relied on decision-making paradigms informed by reinforcement learning models (2, 3) or those involving static, binary choice (4, 5). Recently, however, there has been increased interest across basic (6-8) and clinical (9, 10) science in decision tasks that probe how individuals maximize rewards in more dynamic environments in which the average rate of reward changes over time. In these tasks, individuals make a series of decisions to continue to engage with a particular source of reward that steadily declines in value or leave and search for another, previously unexploited, source. These tasks thus provide an ecologically valid depiction of many real-world decisions and could offer mechanistic insight into the cognitive origins of maladaptive reward pursuit as hypothesized in chronic OUD.

Patch-foraging and the influential Marginal Value Theorem (MVT) (11, 12) are one such class of tasks and models from behavioral ecology used to show how humans and other animals make serial (stay/leave) decisions in a range of contexts, from searching for food to social exchange (6-8). Optimal patch-foraging as prescribed by MVT requires estimating the long-run (average) reward rate of the environment and comparing this estimate to the immediate rate of return at a current reward source (“patch”). This average reward-rate estimate, in turn, depends on the overall (objective) quality of the environment and determines the opportunity cost of time associated with a decision to leave a current resource in pursuit of another (13). Having an accurate estimate of the environmental reward rate is a critical feature of this framework since it can reveal a fundamental computation that renders individuals vulnerable to suboptimal foraging decisions, like leaving patches too early (“underharvesting”) or staying for too long (“overharvesting”) as previously observed under certain conditions like stress (14, 15) and in some neuropsychiatric disorders (10, 16).

Theoretical (17-19) and emerging cross-species empirical work (16, 20-24) has linked decision variables critical to MVT to activity in catecholaminergic systems central to addictive behavior, namely dopamine and norepinephrine. By these accounts, tonic dopamine is suggested to track an estimate of the average reward rate in the environment while tonic norepinephrine modulates decision noise and task disengagement, with increased levels of both promoting earlier patch leaving decisions, albeit through different mechanisms. Both mesolimbic dopaminergic circuits centered on the ventral tegmental area (VTA) and the locus coeruleus (LC) noradrenergic system have long-established roles in addictive behavior in preclinical models (25-30), but their involvement in human OUD remains severely understudied (26, 31). Patch-foraging could thus provide not only a useful translational framework for understanding addiction-relevant behavior but also a window into the pathophysiology of OUD.

Here we used a patch-foraging framework to examine the cognitive mechanisms of chronic opioid use and the putative role of catecholaminergic systems. We examined how the stay/leave decisions of people with chronic OUD are influenced by environmental reward rate, compared to those of matched healthy community controls and the MVT optimal policy. We then tested whether individual differences in foraging behavior could be explained by the function of VTA dopamine and/or LC norepinephrine circuitry using neuromelanin-sensitive (NM-MRI) of the brainstem. Neuromelanin is a product of dopamine – and norepinephrine – metabolism that accumulates with age in the cell bodies of dopamine and norepinephrine producing neurons, where it remains until cell death (32, 33). NM-MRI reliably captures the regional concentration of neuromelanin in catecholaminergic nuclei (34, 35), providing a noninvasive proxy measure of their long-term function (34, 36, 37). A key advantage of this technique is the ability to separately image small nuclei, like the VTA and LC, that are difficult to assess using conventional molecular imaging approaches in humans which have generally low spatial resolution. Given the defining tendency in addiction to pursue the drug despite diminishing returns, we hypothesized that people with OUD would demonstrate overharvesting behavior (i.e., later patch leaving) – relative to healthy comparison controls and the optimal parameters of the MVT. We further hypothesized that overharvesting behavior, indicative of a lower overall estimate of environmental reward rate, would be related to reduced neuromelanin signal in the VTA and/or LC, consistent with potential roles for dopamine and norepinephrine in signaling average reward rate and alterations to their function in human addiction to opioids.

## Results

### Chronic opioid use is associated with a bias to overharvesting in patch-foraging

Participants with moderate to severe OUD (*N*=42) and age, sex, and race/ethnicity matched controls (*N*=33; **Table 1**) completed a patch-foraging task previously used to show that the behavior of healthy adults by and large comports to MVT predictions (13) (**Fig. 1A** and **Methods**). The task consisted of four blocks in which participants made serial decisions to stay and harvest a tree (current patch) for apples worth real monetary rewards, or to exit in search of a replenished tree (new patch) and incur a travel cost (timeout period during which they could not harvest apples for 4.75 s [short blocks] or 10.75 s [long blocks]). The optimal strategy for participants in this task is to track an estimate of the average reward rate in the environment, here differing between blocks only as a function of travel time, and to leave a current tree when its returns drop below this estimate. We denote this value as the *exit threshold*, quantified as the number of apples received prior to an ‘exit’ decision (i.e., at *t*-1) in each tree patch, which we calculated for the MVT optimal case and measured for each participant. **Fig. 1B** shows example participants’ exit thresholds across all patches visited in the task (an average for the group of 48.33±20.68 [*SD*] patches in controls and 37.67±16.25 in OUD).

**Table 1.**
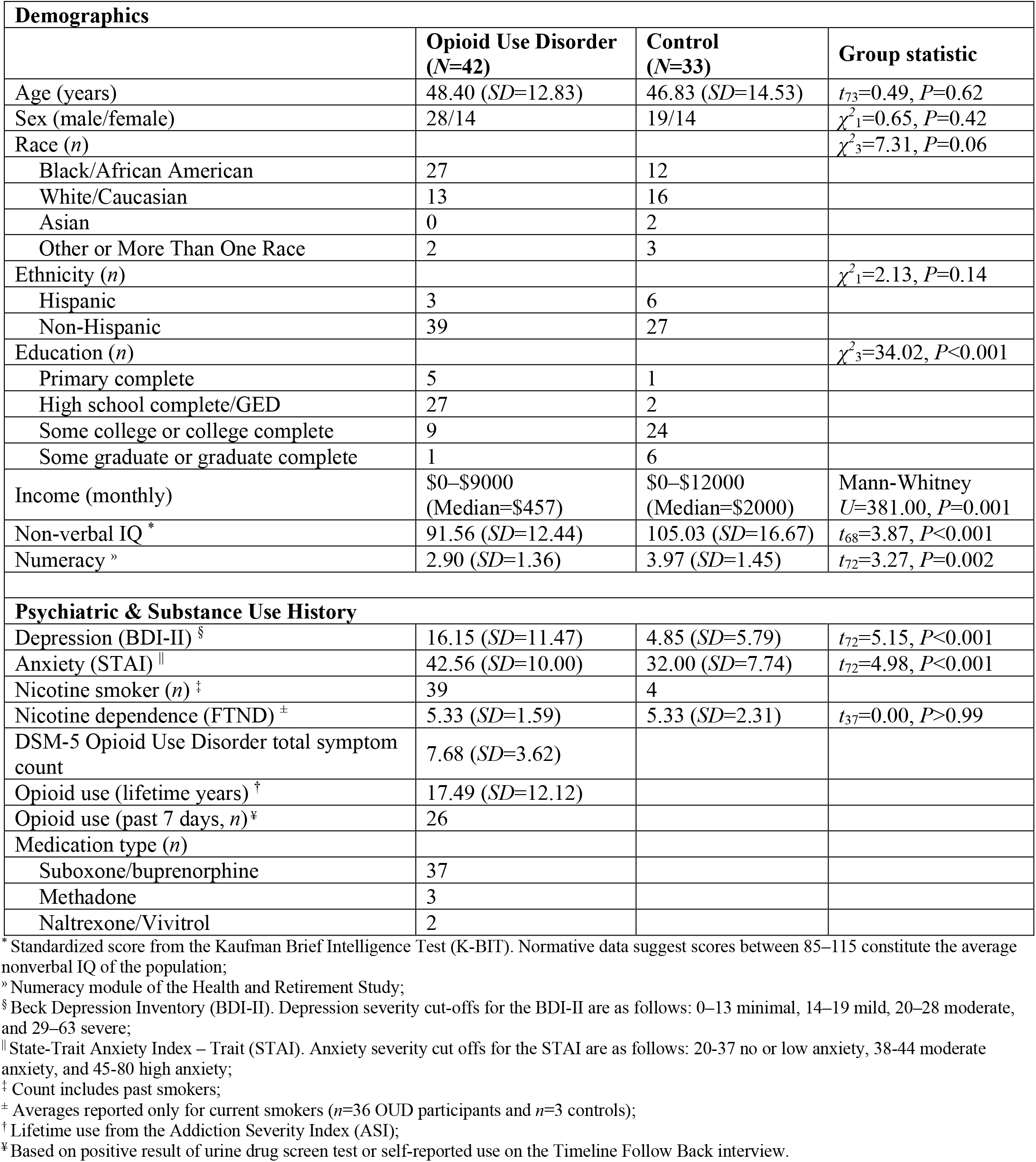
Sample characteristics.

| **Demographics** |  |  |  |
| --- | --- | --- | --- |
|  | **Opioid Use Disorder**<br>**(N=42)** | **Control**<br>**(N=33)** | **Group statistic** |
| Age (years) | 48.40 (*SD*=12.83) | 46.83 (*SD*=14.53) | $t_{73}=0.49, P=0.62$ |
| Sex (male/female) | 28/14 | 19/14 | $\chi^2_1=0.65, P=0.42$ |
| Race (*n*) | | | $\chi^2_3=7.31, P=0.06$ |
| Black/African American | 27 | 12 |  |
| White/Caucasian | 13 | 16 |  |
| Asian | 0 | 2 |  |
| Other or More Than One Race | 2 | 3 |  |
| Ethnicity (*n*) | | | $\chi^2_1=2.13, P=0.14$ |
| Hispanic | 3 | 6 |  |
| Non-Hispanic | 39 | 27 |  |
| Education (*n*) | | | $\chi^2_3=34.02, P<0.001$ |
| Primary complete | 5 | 1 |  |
| High school complete/GED | 27 | 2 |  |
| Some college or college complete | 9 | 24 |  |
| Some graduate or graduate complete | 1 | 6 |  |
| Income (monthly) | \$0–\$9000<br>(Median=\$457) | \$0–\$12000<br>(Median=\$2000) | Mann-Whitney<br>$U=381.00, P=0.001$ |
| Non-verbal IQ * | 91.56 (*SD*=12.44) | 105.03 (*SD*=16.67) | $t_{68}=3.87, P<0.001$ |
| Numeracy » | 2.90 (*SD*=1.36) | 3.97 (*SD*=1.45) | $t_{72}=3.27, P=0.002$ |
| **Psychiatric & Substance Use History** |  |  |  |
| Depression (BDI-II) § | 16.15 (*SD*=11.47) | 4.85 (*SD*=5.79) | $t_{72}=5.15, P<0.001$ |
| Anxiety (STAI) \|\| | 42.56 (*SD*=10.00) | 32.00 (*SD*=7.74) | $t_{72}=4.98, P<0.001$ |
| Nicotine smoker (*n*) ‡ | 39 | 4 |  |
| Nicotine dependence (FTND) ± | 5.33 (*SD*=1.59) | 5.33 (*SD*=2.31) | $t_{37}=0.00, P>0.99$ |
| DSM-5 Opioid Use Disorder total symptom count | 7.68 (*SD*=3.62) |  |  |
| Opioid use (lifetime years) † | 17.49 (*SD*=12.12) |  |  |
| Opioid use (past 7 days, *n*) ¥ | 26 |  |  |
| Medication type (*n*) |  |  |  |
| Suboxone/buprenorphine | 37 |  |  |
| Methadone | 3 |  |  |
| Naltrexone/Vivitrol | 2 |  |  |

\* Standardized score from the Kaufman Brief Intelligence Test (K-BIT). Normative data suggest scores between 85–115 constitute the average nonverbal IQ of the population;
» Numeracy module of the Health and Retirement Study;
§ Beck Depression Inventory (BDI-II). Depression severity cut-offs for the BDI-II are as follows: 0–13 minimal, 14–19 mild, 20–28 moderate, and 29–63 severe;
|| State-Trait Anxiety Index – Trait (STAI). Anxiety severity cut offs for the STAI are as follows: 20-37 no or low anxiety, 38-44 moderate anxiety, and 45-80 high anxiety;
‡ Count includes past smokers;
± Averages reported only for current smokers ( $n=36$ OUD participants and $n=3$ controls);
† Lifetime use from the Addiction Severity Index (ASI);
¥ Based on positive result of urine drug screen test or self-reported use on the Timeline Follow Back interview.

**Fig. 1.**
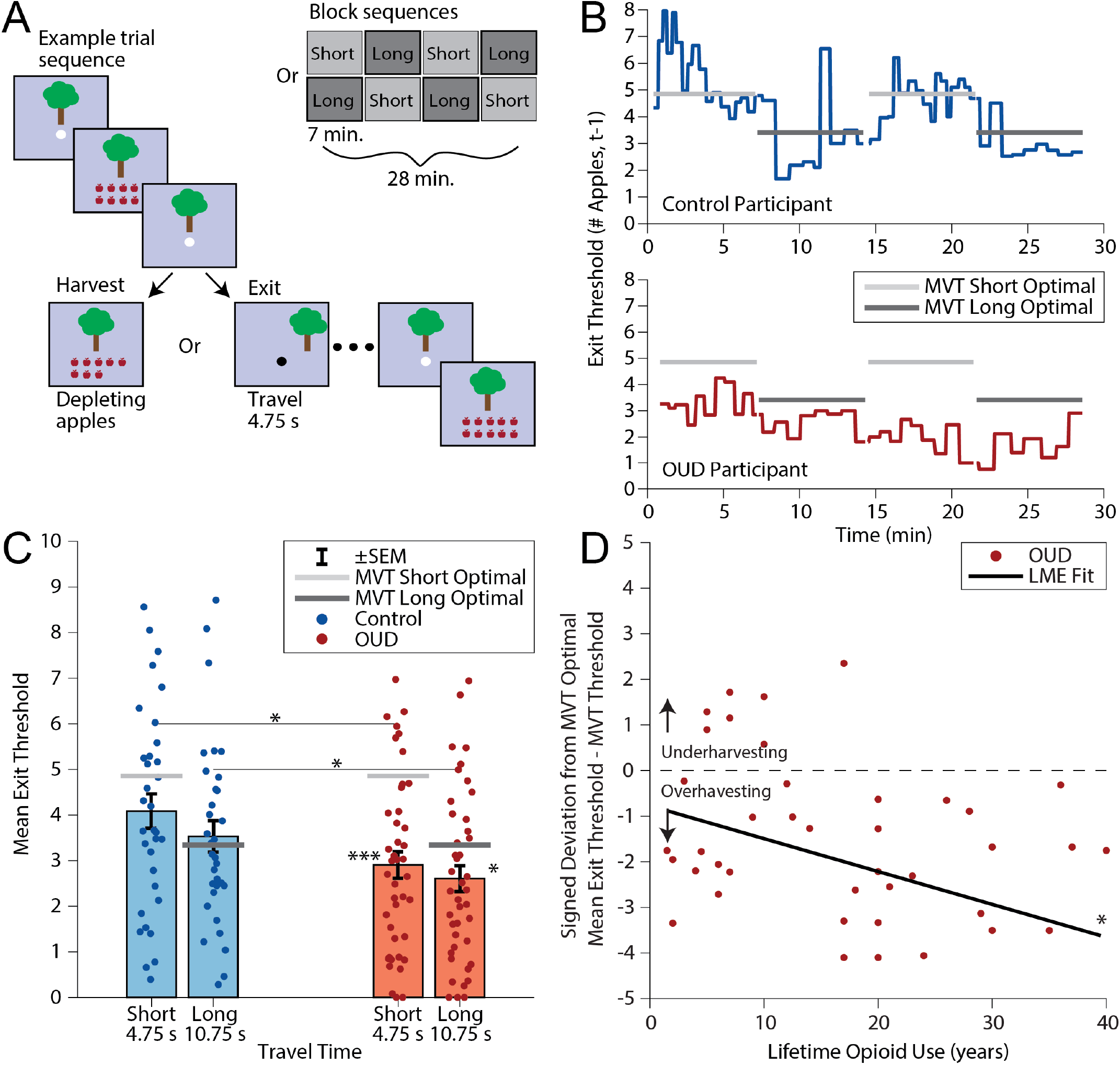
Foraging task and behavior. (**A**) Participants completed 4 blocks of the foraging task, with blocks having either short (4.75 s) or long (10.75 s) travel times between trees. Blocks were counterbalanced across participants (ABAB or BABA order) and signaled by a change in background color. On each trial within a block, participants made serial decisions to harvest a current tree for apples (to be converted to money at the end of the task) or to travel to a new, replenished tree but incur a travel time cost. The number of apples (rewards) received from a given tree decayed exponentially with each harvest decision. If participants chose to exit and travel to a new tree, they had to wait to collect additional apples for a timeout period equal to the travel time in the current block. (**B**) Illustrative examples of trial-by-trial exit thresholds (number of apples received prior to an exit decision, i.e., at *t*-1) from a control participant and a participant with opioid use disorder (OUD) across the entire task, plotted against the optimal threshold as predicted by the Marginal Value Theorem (MVT). (**C**) Mean exit thresholds by diagnostic group and travel time, showing lower exit thresholds in participants with OUD (indexed by lower values) relative to controls and MVT predicted thresholds. By contrast, controls do not differ significantly from optimal at either short or long travel time. (**D**) Lower exit thresholds in participants with OUD (indexed by lower values) correlate with longer duration of lifetime opioid use, controlling for age. MVT, marginal value theorem; OUD, opioid use disorder; SEM, standard error of the mean. * *P*<0.05, *** *P*<0.001.

Across the entire sample, trial-by-trial exit thresholds were higher in the short travel time blocks than the long travel time blocks (short, long; *B*=-0.09, 95% *CI* [-0.15, -0.04], *t*_66.77_=-3.65, *P*=0.0005) and in controls relative to participants with OUD (*B*=-1.38, 95% *CI* [-2.47, -0.29], *t*_74.10_=-2.52, *P*=0.01), but there was no significant interaction between travel time and diagnosis (*B*=0.04, 95% *CI* [-0.03, 0.11], *t*_69.55_=1.22, *P*=0.23; see **Methods**). Thus, as expected—and confirming exit thresholds reflect internal estimates of the different average reward rates of the two environments—participants had higher exit thresholds (left a patch sooner) when the travel time to a newly replenished patch was shorter. Critically, while OUD participants did not differ significantly in their sensitivity to travel time, they had markedly lower exit thresholds overall (left patches later) than controls (**Fig. 1C**). Analyses of stay/leave decisions as an alternative behavioral measure are reported in **SI Results**; these led to the same conclusion.

We next compared participants’ exit thresholds against the reward-maximizing (optimal) strategy given by the MVT. While controls’ exit thresholds did not significantly deviate from MVT optimal thresholds, we observed that OUD participants harvested on average 1.95±0.29 apples more than optimal before exiting a tree in short blocks, and 0.74±0.28 apples more than optimal in long blocks. This led to about 25% and 8% less in task earnings, respectively, relative to the MVT reward-maximizing strategy and to controls (controls: $24.02±3.66, OUD: $22.11±4.46; *t*_73_=-1.99, *P*=0.049).

To formally assess for differences in over/underharvesting, we repeated our group analysis after first subtracting the MVT optimal threshold for each block from participants’ trial-by-trial exit thresholds. Results provided clear support for overharvesting: signed deviations from MVT were negative and bigger for the short travel time blocks (*B*=0.16, 95% *CI* [0.11, 0.21], *t*_66.77_=6.14, *P*<0.0001) and overall in the OUD group (*B*=-1.38, 95% *CI* [-2.47, -0.29], *t*_74.10_=-2.52, *P*=0.01; interaction effect: *P*=0.23). Suggestive of a direct relationship to drug use, we further found that overharvesting worsened with increased lifetime opioid use in those with OUD (controlling for age, *B*=-0.10, 95% *CI* [-0.17, -0.03], *t*_46.35_=-2.85, *P*=0.006, two influential outliers excluded, Cook’s *d*>0.07; **Fig. 1D**). Again, there was no significant interaction with travel time (*P*=0.11). By contrast, sociodemographic factors that differed between groups, such as income and IQ, were unrelated to overharvesting (**Table 1, SI Results**). In support of overharvesting being driven primarily by differences in internal estimates of the environmental reward rate rather than lower-level factors such as task/attentional disengagement, task disengagement as indexed by decision noise and the frequency late-responses was generally low and comparable between groups (**SI Results**). Thus, as previously observed in healthy young adults (13), while participants adjusted their behavior to changes in the long-run reward rate of the environment, they systematically deviated from the reward-maximizing strategy, and this was especially pronounced in more chronic OUD participants who exhibited a strong tendency to overharvest.

### Overharvesting selectively relates to NM-MRI signal in dopaminergic nuclei

To assess catecholaminergic contributions to overharvesting, in a subset of participants (*n*=25 OUD and *n*=28 controls), we collected NM-MRI scans of the brainstem with submillimeter in-plane resolution. For each participant, we computed the contrast ratio for each voxel in the preprocessed images relative to signal in a control white-matter region with negligible neuromelanin content, the crus cerebri (38) (see **Methods**). The resulting contrast maps show regions with high neuromelanin accumulation as hyperintense, with interindividual variability in NM-MRI signal contrast indexing differences in neuromelanin concentration in these regions and, indirectly, the long-term function of dopamine and norepinephrine neurons therein (32-37). Our analyses focused on the VTA and LC as *a priori* regions of interest, given suggested dopaminergic and noradrenergic involvement in foraging (16-24) and preclinical evidence for VTA and LC dysregulation with opioid exposure (25-30). **Fig. 2** shows the averaged NM-MRI contrast map of the sample (**A** and **B**, left) with the region-of-interest masks overlaid (right), confirming good spatial coverage of both regions.

**Fig. 2.**
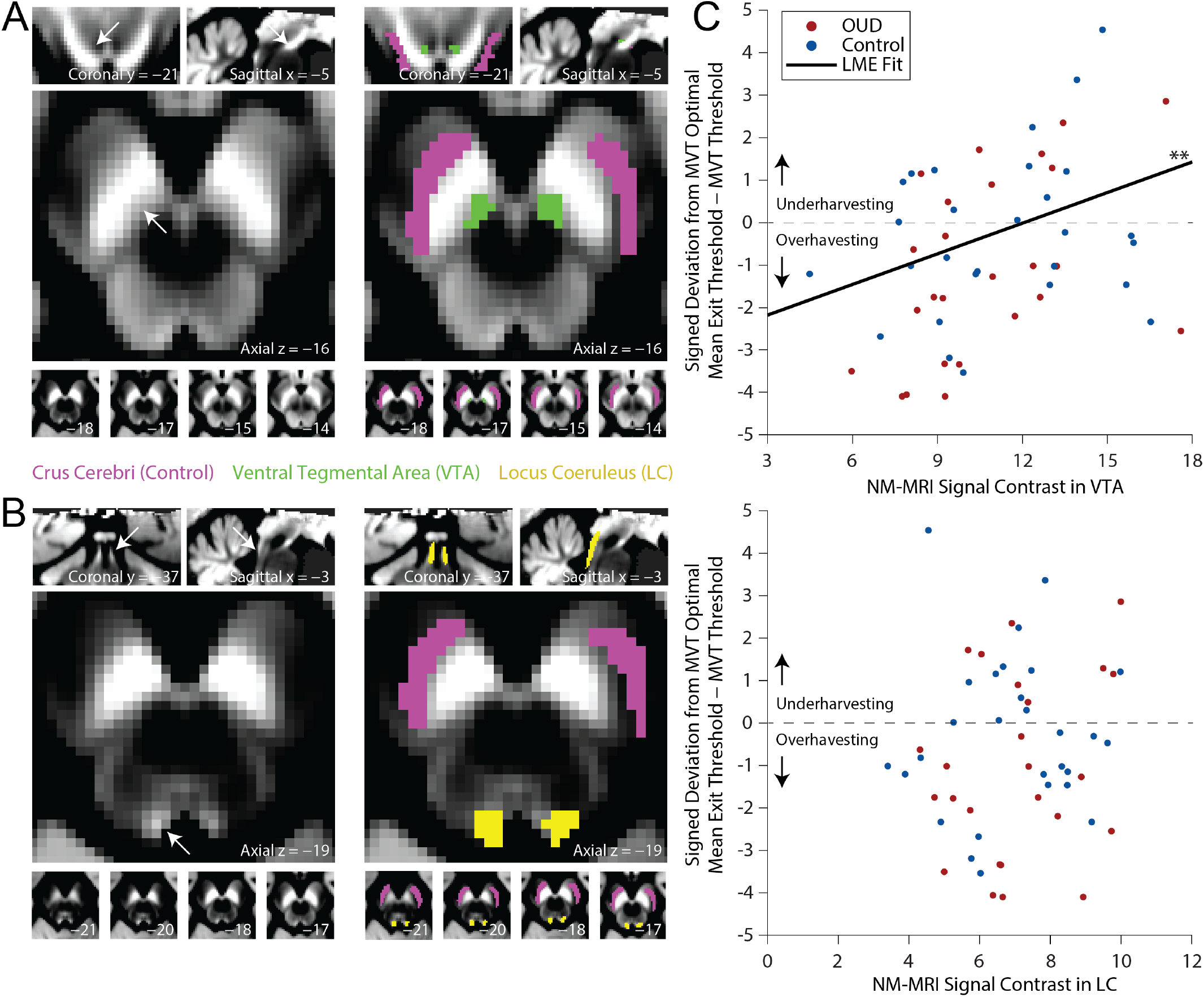
Region-of-interest analyses of the relationship between neuromelanin signal and foraging behavior. Group average contrast ratio map showing neuromelanin signal in each voxel of the preprocessed NM-MRI scans relative to a control region (the crus cerebri, CC), at the location of (**A**) the ventral tegmental area (VTA; left) with a probabilistic VTA mask (39) overlaid at a threshold of 0.5 (right; displayed at MNI coordinates *x*=-5, *y*=-21, *z*=-16) and (**B**) the locus coeruleus (LC; left) with a probabilistic LC mask (40) overlaid at a threshold of 0.05 (right; displayed at MNI coordinates *x*=-3, *y*=-37, *z*=-19). (**C**) Increased neuromelanin signal contrast in the VTA correlates across all participants with less overharvesting (signed deviation of participants’ exit thresholds from the MVT optimal threshold, controlling for age and repetition time [TR] of the scan acquisition). No significant relationship is observed with LC neuromelanin signal contrast. CC, crus cerebri, LC, locus coeruleus; MVT, marginal value theorem; OUD, opioid use disorder; VTA, ventral tegmental area. ** *P*<0.01.

Using average signal in the two regions of interest as predictors of trial-by-trial signed deviation of participants’ exit thresholds from the MVT optimal thresholds (our measure of over/underharvesting) revealed only a significant effect for the VTA (*B=*0.24, 95% *CI* [0.07, 0.41], *t*_53.07_=2.77, *P=*0.008) but not the LC (*B=*0.07, 95% *CI* [-0.29, 0.42], *t*_52.88_=0.38, *P=*0.70), such that individuals with higher neuromelanin signal contrast in the VTA, but not in the LC, exhibited less overharvesting behavior (**Fig. 2C** and **Table S1**). No significant interaction effects were observed with diagnosis or travel time when these were included as additional predictors in the models (*P>*0.19), and the same analyses only within the OUD group revealed the same selective VTA effect (*P=*0.006). We also evaluated the relationship between VTA and LC signal contrast and task/attentional disengagement and found no effects (**SI Results**). This supports a relationship between reduced VTA neuromelanin and overharvesting that is present across the entire sample and within those with OUD specifically, and that is indicative of lower perceived environmental reward rate but not increased task/attentional disengagement.

### Exploratory analyses of subregional effects confirm specificity to ventral tegmental area

To further characterize the anatomy of dopaminergic subregions relevant to overharvesting behavior, in a region-of-interest agnostic manner, we examined the distribution of voxels within the broader substantia nigra (SN)/VTA complex that showed a positive relationship with behavior. Strikingly, the voxels showing this relationship (171/1,345 at a α<0.05 threshold) included almost the entirety of the VTA (**Fig. 3A**). Partitioning the SN/VTA complex into the SN pars compacta, SN pars reticulata, and VTA subregions using a probabilistic atlas (39) indicated 79% of the total number of voxels in the VTA, and only 8% and 3% of those in the SN pars compacta and SN pars reticulata, respectively, correlated with less overharvesting (**Fig. 3B**; *χ*^*2*^_2_=246.99, *P*<1.0×10^−307^). These results were robust to different mask probability thresholds and alternative anatomical definitions of the VTA that include the parabrachial nucleus (41), further showing that voxels more strongly related to overharvesting had higher probability of belonging to the VTA (**Fig. S1** in **SI Results**). Collectively, these findings indicate that a bias to overharvest selectively relates to putative dopaminergic function in the VTA.

**Fig. 3.**
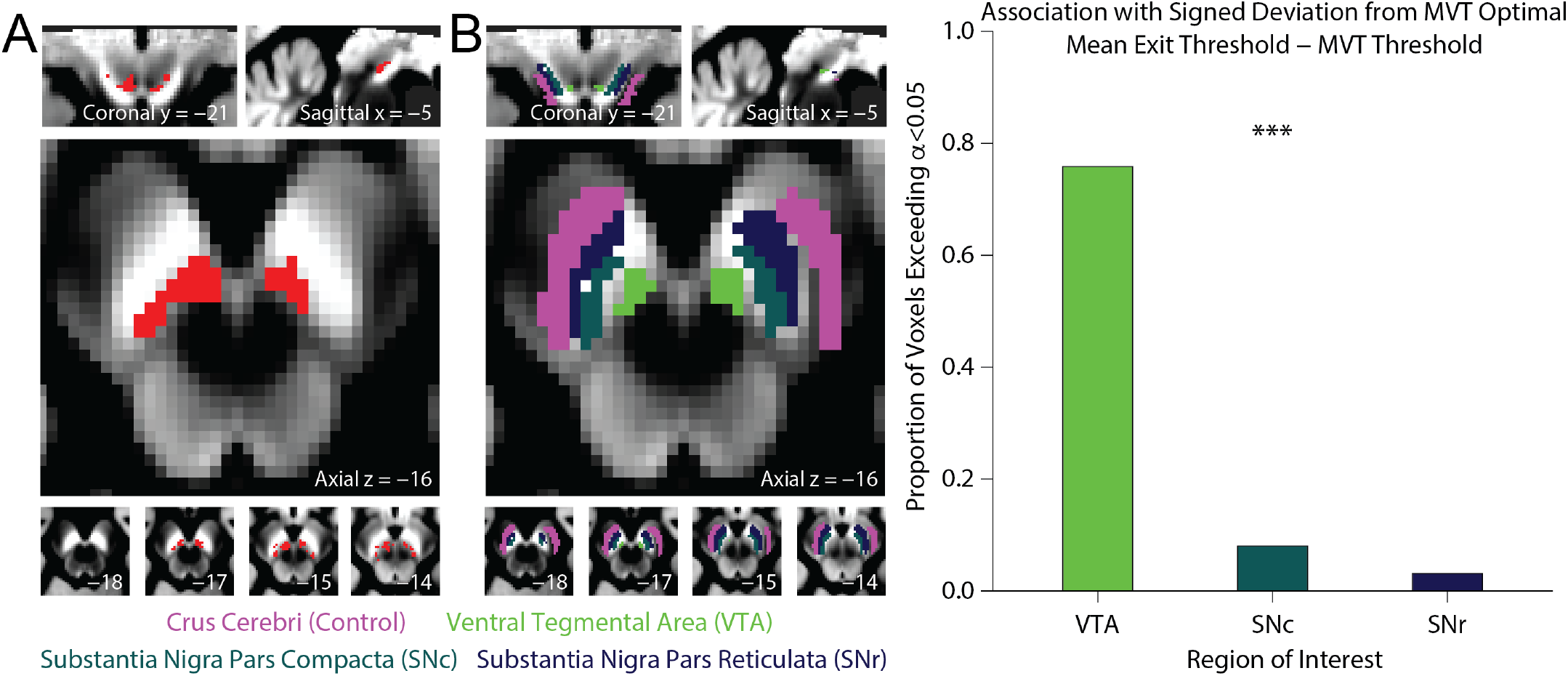
Voxel-wise analysis of the relationship between neuromelanin signal in dopaminergic subregions and foraging behavior. Group average contrast ratio map showing neuromelanin signal in each voxel of the preprocessed NM-MRI scans relative to a control region (the crus cerebri, CC), showing (**A**) voxels (in red) exceeding α<0.05, uncorrected relationship to less overharvesting behavior in voxel-wise analyses within a mask of the broader substantia nigra/ventral tegmental area (SN/VTA) complex (controlling for age and repetition time [TR] of the scan acquisition; displayed at MNI coordinates *x*=-5, *y*=-21, *z*=-16). (**B**) Quantification of the proportion of total voxels within the VTA, SN pars compacta (SNc), and SN pars reticulata (SNr) in which increased neuromelanin signal correlates with less overharvesting (at α<0.05), confirming this relationship is highly selective to the VTA. CC, crus cerebri; MVT, marginal value theorem; OUD, opioid use disorder; SN, substantia nigra; VTA, ventral tegmental area. *** *P*<0.001.

Further supporting this conclusion, a *post hoc* region-of-interest analysis including all three dopaminergic subregions (VTA, SN pars compacta, SN pars reticulata) and the noradrenergic LC in the same model as predictors of participants’ behavior revealed only a significant relationship between neuromelanin signal in the VTA and less overharvesting (*B*=0.48, 95% *CI* [0.16, 0.79], *t*_53.45_=3.02, *P*=0.004; **Table S2**). A similar analysis predicting years of opioid use and, separately, OUD diagnosis also provided preliminary evidence for selectivity to the VTA: only reduced VTA neuromelanin signal was a significant, unique, predictor of both longer use (*B=-*2.85, *t*_18_=-2.17, *P*=0.04; **Table S3**) and increased odds of belonging to the OUD group versus the control group (*B=-*0.53, *t*_46_=-2.08, *P*=0.04; **Table S4**), pointing to a shared neurobiological mechanism of overharvesting in patch-foraging decisions and human opioid addiction.

## Discussion

We used a patch-foraging framework combined with NM-MRI to test how a behavior purportedly associated with maladaptive reward pursuit relates to catecholaminergic systems implicated in addiction. We found that adults with OUD exhibited a strong tendency to overharvest relative to matched healthy controls and an optimal policy informed by the MVT. This tendency was most pronounced in those with more chronic opioid use histories. We then examined whether interindividual variability in overharvesting related to neuromelanin signal, and indirectly catecholaminergic function, of the VTA and LC. We found that individuals with increased neuromelanin signal in the VTA overharvested less but did not observe an association with the LC. The correlation with VTA signal was robust and selective to the VTA compared to other dopaminergic nuclei. These findings provide support for alterations in patch-foraging (serial stay/leave) decisions in the pathophysiology of human OUD and implicate the VTA in this process, in line with an extensive preclinical literature highlighting a central role for VTA-dependent circuitry in addiction.

Optimal patch-foraging in the wild requires maintaining a dynamic estimate of the average reward rate in the environment and leaving a current course of action when its immediate rate of return falls below this estimate. To mimic these conditions in the lab, participants completed a virtual patch-foraging task where they harvested apples to earn monetary rewards. Critically, these decisions were made in two environments that differed only in their travel time requirement and thus average reward rate. Both groups adjusted their behavior to changes in the environmental reward rate, leaving patches sooner when the average reward rate was higher and staying longer when it was lower. However, individuals with OUD, and especially those with more chronic histories of opioid use, showed a consistent tendency to stay longer in a patch relative to controls and the MVT optimal strategy, which led to overall lower task earnings.

Our data suggest that this tendency to overharvest in the OUD group may be explained by biased estimates of reward rate, and not decision noise or a global learning deficit. Decision noise such as variability in exit thresholds and reaction times was unrelated to diagnosis or years of opioid use. The requirement for learning was minimized by varying only travel time between blocks and by providing participants with explicit instructions on the task structure (and indeed, both groups showed the expected sensitivity to travel time). This also points to an important distinction between our patch-foraging task and other explore/exploit paradigms in which participants decide between choosing options with known reward values vs. exploring the environment for options that may yield greater rewards. While explore/exploit trade-offs may ostensibly appear similar to the patch-foraging approach used here, a critical difference is that participants are fully aware that replenished patches are available whenever they choose to leave, thus eliminating the uncertainty characteristic of exploratory decisions in these standard explore/exploit tasks. We also note that while other factors such as risk aversion and temporal discounting, which have previously been found to differ in OUD (42-53), could contribute to foraging decisions, their influence in the current study is likely small. Later patch-leaving would be predicted by increased risk aversion; however, people with OUD have been found to be more risk tolerant (47-53), which would predict underharvesting rather than the overharvesting behavior we observed. Similarly, people with OUD have steeper temporal discounting (42-46). Prior work however suggests that a temporal difference learning model of the current task that incorporates a discounting factor cannot fully explain later patch-leaving and generally provides a poorer behavioral fit than the MVT model (13). And while increased hyperbolic discounting does predict staying in patches longer overall, in our task it also predicts increased sensitivity to travel time which we did not find in OUD. Thus, the overharvesting bias observed in OUD participants is most parsimoniously explained by a misestimation of environmental reward rate, providing an intuitive explanation for individual differences in reward pursuit despite diminishing returns and when alternative sources of reward are available, as observed clinically in the chronic stages of OUD.

Distinct from phasic dopamine responses critical for prediction error encoding, emerging findings suggest tonic dopamine provides a neural mechanism for tracking of a long-run estimate of environmental reward rates necessary for optimal patch-foraging (16, 17, 19-21, 23, 24). Here we show that overharvesting relates to individual differences in VTA neuromelanin signal, a proxy measure of long-term dopamine function that is likely influenced by slower dopamine changes. These data directly implicate mesolimbic dopamine systems in patch-foraging and provide *in vivo* evidence for VTA alterations in human addiction to opioids. Dopamine hypofunction has been consistently observed in animal models of OUD where the long-term effects of opioids result in inhibition of dopamine neurons in the VTA via their action on tonically active GABAergic interneurons (26, 27, 54). This results in VTA-accumbens circuit disruption and lowered dopamine tone. While molecular PET studies in humans have consistently found dopaminergic disruption across addiction disorders (30, 55-57), studies of OUD have been scarce (58), and anatomical specificity to the VTA has been challenging to demonstrate. Like rodents chronically exposed to nicotine who develop alterations in tonic midbrain dopamine firing rates and adopt exploitative decision strategies (59), here we find that people with chronic OUD exhibit a tendency to overharvest which correlates selectively with reduced neuromelanin signal in the VTA. These findings extend a large body of animal work on chronic opioid use to humans and provide translational evidence that mesolimbic dopamine circuits underlie drug-relevant reward behaviors and thus constitute candidate targets for therapeutic intervention in OUD.

Given established alterations in both VTA dopamine and LC norepinephrine circuits in preclinical models of OUD (25-30), and that both increased tonic dopamine and norepinephrine could lead to earlier patch-leaving (16-24)—albeit through different mechanisms—we hypothesized a potential role for both catecholamines in the current study. While our imaging protocol was optimized to separately capture both nuclei, we only found evidence that VTA neuromelanin signal correlated with foraging behavior. Given LC involvement in foraging may be less straightforward, involving changes in its phasic and tonic modes, it is possible that NM-MRI may not be well-suited to capture this complexity. However, prior work using NM-MRI of the LC has found broad cognitive correlates related to attention and memory function (60, 61), suggesting this measure may index interindividual variability in norepinephrine for tasks relying on distinct cognitive processes than those engaged here. Prior work in human cocaine addiction has also found evidence for increased LC neuromelanin signal (62). No diagnostic group differences were reported in the VTA in that study or in another focused on the substantia nigra which also found increased substantia nigra neuromelanin signal (63). Thus, the specific conditions in which LC norepinephrine function as captured by NM-MRI contributes to addictive behavior remains to be determined, as well as the impact of different drugs of abuse on these circuits.

While prior work indicates NM-MRI serves as a reliable proxy measure of the function of catecholamines (38) and shows promise as a clinical marker of disorders of these systems even in the absence of neurodegeneration, including depression and psychosis (64-66), the precise relationship to tonic (vs. phasic) dopamine remains unclear. Studies show chronic L-DOPA, which is thought to increase both phasic and tonic dopamine, increases neuromelanin concentration in both humans and rodents (66, 67), providing some assurance that NM-MRI could serve as a reasonable proxy of these neurobiological processes. However, further work is needed in this area, including in the context of OUD. In addition, while we found that overharvesting increased with longer history of opioid use, participants in the current study were also receiving opioid substitution medications and the combined impact of these medications and illicit opioids is unknown. Nevertheless, both types of opioid exposure are likely reflections of the chronicity and severity of the disorder. Future work is needed to explore how standard treatment for OUD affects patch-foraging and its underlying neurobiology. NM-MRI can also reflect changes in cell density or morphology, but it is unlikely that OUD participants have substantial dopamine cell loss, and even though morphological changes in VTA neurons are possible (68), these may be restricted to certain cell types (69) and are in any case linked to functional changes (68).

In summary, we find that foraging behavior in human addiction relates to the long-term function of VTA dopamine systems that have long been hypothesized to mediate this class of ecologically valid stay/leave decisions. Our results demonstrate maladaptive reward pursuit in OUD may be driven in part by misestimation of environmental reward rate, which has implications for continued drug-seeking behavior despite diminishing returns characteristic of this disorder, thereby advancing a translational framework that links VTA-dependent circuitry to drug addiction.

## Methods

### Participants

OUD participants were recruited from university-affiliated outpatient medications for OUD programs. Controls were recruited from the same geographic area to be matched to patients on age, sex, and race/ethnicity. Inclusion criteria for both groups were ≥18 years of age, ability to provide informed consent, and ability to understand and complete study procedures. Participants were included in the OUD group if they had a primary diagnosis of OUD encompassing heroin and/or painkiller use, as obtained from patient charts, had ≥12 months history of opioid use and were currently treatment-engaged. Exclusion criteria for both groups were: active psychosis or mania; current or past diagnosis of schizophrenia; history of intellectual disability or developmental or neurological disorder; history of seizures or epilepsy; history of loss consciousness >30 minutes; severe medical conditions requiring hospitalization or that could compromise study participation (e.g., renal or liver failure, end-stage AIDS); and for those willing to participate in the MRI component, MRI contraindications including metal in the body and pregnancy. Control participants were further excluded if they had: a positive urine drug screen on any study day; current or past problematic substance use other than nicotine and alcohol abuse confined to college or military service; or current or past bipolar disorder diagnosis. Informed consent to participate was obtained in accordance with procedures approved by the Rutgers University IRB.

A total of *n*=3 participants across both groups were excluded *post hoc* for failing to meet inclusion/exclusion criteria; in addition, *n*=2 controls were excluded for being closely genetically related and *n*=3 for being a poor sociodemographic match. Following these exclusions, *N*=42 OUD participants and *N*=33 controls were included in the analyses reported. The groups were matched on age, sex, race, and ethnicity, however the OUD group had significantly lower educational achievement, non-verbal IQ, and numeracy, and significantly higher severity of depression and anxiety relative to controls (**Table 1**). Most OUD participants were receiving suboxone/buprenorphine and had used illicit opioids in the previous month. All participants completed screening procedures and the foraging task; a subset (*n*=53) of eligible and willing participants also completed the MRI procedures. Payment for participation was $10/h plus a task performance bonus (see below), and an additional $30 for the MRI.

### Foraging task

The task was a variant of Experiment 1A in Constantino et al. (13). In each trial, participants made serial decisions to stay in a current tree patch and harvest rewards or to leave in search of a new one (**Fig. 1A**). Rewards were depicted by apples, which were converted money at a fixed exchange rate of one whole apple to 1 cent and paid out in total to participants at the end of the task as a performance bonus. Participants indicated their choice to stay or leave by one of two key presses when prompted by a response cue (down or right arrow). With each decision to stay and harvest apples (i.e., rewards), the apple supply declined according to a randomly drawn multiplicative factor, such that on average the obtained rewards decayed exponentially. If the participant decided to leave for a new, replenished tree (exit decision), they incurred a travel time delay. Upon arriving to a new patch, participants again needed to decide whether to stay and harvest the current patch or leave and explore other patches. The total task consisted of 4 blocks with a fixed duration of 7 min (or 28 min in total). Blocks were counterbalanced in ABAB/BABA block order across participants and differed only in the travel time between trees (i.e., short=4.75 s, long=10.75 s). New blocks were signaled by a change in background color. The reaction time of each choice was counted toward the ensuing harvest or travel delay so that the total interval between response cues (and thus the average reward rate) was unaffected by response speed.

We varied travel time across blocks to create high and low average reward rate foraging environments; all other features of the task environment were kept constant. The initial supply of apples for each tree was randomly drawn from a Gaussian distribution with a mean of 10 (*SD*=1.00). The depletion rate for each successive harvest of a tree was drawn from a Beta distribution with parameters α=14.91 and β=2.03. These parameters were set so the mean rate of depletion was 0.90 (*SD*=0.09). Participants were informed of the total task length, that travel times could vary across blocks (but not within a given block), and that trees would vary in their quality and depletion rate. Participants completed a practice block to get familiar with the contingencies of the task in which earnings did not count toward the bonus.

### Image acquisition and preprocessing

All scanning was performed on a 3T Siemens TRIO using a 32-channel head coil. We first acquired a high-resolution T1-weighted image using a 3D magnetization prepared rapid acquisition gradient echo (T1w MPRAGE) sequence with the following parameters: spatial resolution=0.8×0.8×0.8 mm; field-of-view (FOV) read=256 mm; 208 slices; echo time (TE)=2.31 ms; repetition time (TR)=2400 ms; flip angle=8°; in-plane acceleration GRAPPA=2; bandwidth=210 Hz/pixel. Neuromelanin-sensitive MRI scans were acquired with a 2D turbo spin echo (TSE) sequence with the following parameters: in-plane (spatial) resolution=0.6875×0.6875 mm; slice thickness=1.5 mm, FOV=220 mm^2^; 20 slices; TE=12.2 ms; TR^a^=633 ms; flip angle=120°; bandwidth=180 Hz/pixel; and 7 averages. The slice-prescription protocol consisted of orienting the image stack along the AC-PC line using each participant’s T1-weighted scan as reference and placing the top slice 10.5 mm above the top of the pons, as viewed on the most medial slice in the sagittal plane. This protocol ensured coverage of the entire LC, the entire VTA, and most of the substantia nigra except for its most dorsal aspects in some individuals.

The NM-MRI data were preprocessed using an optimized pipeline (70) based on ANTs routines (71, 72) to allow for analyses in standardized MNI space. This included: 1) brain extraction of T1w images using “antsBrainExtraction.sh”; 2) spatial normalization of the brain-extracted T1w images to the MNI152NLin2009cAsym template space using “antsRegistrationSyN.sh”; 3) co-registration of the NM-MRI images to the T1w images using “antsRegistrationSyN.sh” (rigid); 4) spatial normalization of the NM-MRI images to template space by a single-step transformation combining the transformations estimated in steps 2 and 3 using “antsApplyTransforms.sh”; and 5) spatial smoothing of the normalized NM-MRI images with a 1 mm FWHM Gaussian kernel bounded by the brain mask estimated in step 1 using AFNI’s “3dBlurInMask”. All preprocessed data were individually visually inspected for quality control purposes.

For each participant, a map of contrast-ratios at each voxel *v* was calculated as the relative difference in NM-MRI signal intensity *I* from a reference region *RR* of white matter tracts known to have minimal neuromelanin content, the crus cerebri (38), as follows:

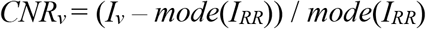

A template mask of the reference region was created by manual tracing in MNI space on an average of normalized NM-MRI scans from all participants in an independent sample [see (70) for more details]. The *mode(I*_*RR*_) was calculated for each participant from a kernel-smoothing-function fitted to a histogram of the distribution of all voxels in the mask.

### Statistical analyses

We first computed the MVT reward-maximizing exit threshold value for each block type (short and long travel time) based on the long-run average reward rate implied by the parameters of the environment, as described in (13). These values were 3.341 for long blocks and 4.855 for short blocks. We then compared participants’ trial-by-trial exit thresholds between travel time blocks and diagnostic groups, and against the MVT values. Lastly, we examined individual differences within the OUD group with respect to years of opioid use, and within all participants, with respect to neuromelanin signal contrast in key regions of interest. Age was included as a covariate in these analyses given its relationship to both years of use and neuromelanin accumulation (32, 73, 74). TR was also included as a covariate in all neuroimaging analyses.

Our primary analytic approach was linear mixed-effects regression, with exit threshold (or deviation of the exit threshold from the MVT optimal threshold) for participant *i* at trial *t* as the outcome variable. The initial behavioral model included predictors for travel time (coded on a numeric scale: 4.75 [short] or 10.75 [long]), diagnosis (coded as a factor: OUD or control), and their interaction. The same model in OUD participants included predictors for years of use and age instead of diagnosis. The initial neuroimaging models included predictors for mean signal contrast in the bilateral VTA and LC, age, and TR of the scan acquisition. Extended models included mean signal in the SN pars compacta and SN pars reticulata as additional predictors. All models were estimated in MATLAB using *fitlme* and included random intercepts and random slopes for travel time by participant. Degrees of freedom for significance testing were computed using Satterthwaite approximation.

For neuroimaging analyses, the mean signal contrast for each region of interest was extracted from *a priori* defined masks (39, 75), excluding voxels with a contrast ratio<0. For the VTA and SN subregion probabilistic masks (39), we used a conservative probability threshold of 0.5 to avoid regional cross-contamination and because different dopaminergic nuclei are expected to have different functional roles in motivation and cognition (75). For the LC (40), we used a more lenient probability threshold of 0.05 to account for its small size and to ensure LC inclusion despite interindividual variability in anatomy (confirmed through careful visual inspection) and because cross-contamination of adjacent catecholaminergic structures is not a concern for the LC. We note however that all results were robust to mask threshold (**SI Results**). Our primary approach to the neuroimaging data was region-of-interest analysis of neuromelanin signal contrast in the VTA and LC. Given the small size of these regions, we deemed voxel-wise tests focused on the spatial extent of effects unsuitable to test the *a priori* hypotheses. In *post hoc* analyses of voxel-wise effect patterns, we determined the subregional mapping of voxels in which neuromelanin signal contrast correlated with participants’ behavior within the entire bilateral SN/VTA complex (agnostic to subregion). Voxels that had a contrast ratio value more extreme than the 1^st^ and 99^th^ percentile across all participants were censored. Voxels correlated with behavior (deviation in exit thresholds from MVT) at α<0.05, uncorrected, were mapped back to region-of-interest demarcations to determine their subregional specificity.

## Acknowledgements

This work was supported by grants from the National Institute on Drug Abuse (R01DA053282 and R01DA054201 to A.B.K.) and the National Institute of Mental Health (R01MH117323 and R01MH114965 to G.H.), a Busch Biomedical Research Grant (to A.B.K.), and a Rutgers Center for Alcohol and Substance Use Studies Pilot Award (to K.B.). We would like to thank Julia Kong and Sahar Hafezi for help with data collection, Cliff Cassidy for help with NM-MRI sequence optimization, and all the patients and staff at the Rutgers SATS and CARE Outpatient Treatment Programs.

## Author Contributions

C.M.R., S.M.C., G.H, and A.B.K. designed the study. K.B., D.B., and J.X. collected the data. A.K. and K.W. analyzed the data. C.M.R., K.B., A.K., and A.B.K. wrote the first draft of the manuscript. All authors provided comments on the final version.

## Footnotes

a For some participants, a SAR warning required increasing the TR. For these participants, TRs ranged from 650-1070 and as a result led to longer scan times. In all NM-MRI analyses, TR was included as a covariate; none of the results reported were affected by TR length.

