## Supplemental Information for "Suboptimal foraging decisions and involvement of the ventral tegmental area in human opioid addiction"

#### Supplemental Results

*Behavioral results: testing for potential sociodemographic confounds*

To test whether sociodemographic factors that differed between the diagnostic groups (see **Table 1**) could potentially confound the identified group difference in foraging, we performed a backwards-elimination stepwise analysis using the sociodemographic factors as predictors of participants' exit thresholds (*stepwiselm* in MATLAB). The only variables to survive the backwards-elimination procedure were nonverbal IQ (KBIT total score) and monthly income. In a subsequent linear mixed effects analysis predicting signed deviation of participants' exit thresholds from the MVT optimal threshold from nonverbal IQ score, monthly income, and travel time as fixed effects, with random slopes and intercepts by participant, we found that monthly income was a significant predictor of earlier patch leaving ($B$=0.0002, 95% $CI$ [$4.67\times10^{-06}$, 0.0004], $t_{69.59}$=2.05, $P$=0.045) while nonverbal IQ was not ($P$=0.97). However, when diagnosis was introduced in the model, we continued to observe a significant diagnosis effect ($B$=-1.21, 95% $CI$ [-2.34, -0.09], $t_{77.59}$=-2.14, $P$=0.035) while the monthly income effect became nonsignificant ($P$=0.28), suggesting that differences in monthly income (or any of the other sociodemographic variables considered) did not explain differences in foraging behavior observed between participants with opioid use disorder (OUD) and controls.

*Behavioral results: trial-by-trial stay/leave decisions*

As a complementary analysis to using exit thresholds as our primary behavioral outcome, we also performed a generalized linear mixed effects analysis predicting the probability of exiting the current patch at each trial in the task (1=exit, 0=harvest). Using travel time, diagnosis and random intercepts and slopes by participant as predictors, we found significant travel time ($B$=-0.02, 95% $CI$ [-0.04, -0.007], $t_{33,745}$=-2.74, $P$=0.006) and diagnosis ($B$=-0.39, 95% $CI$ [-0.71, -0.08], $t_{33,745}$=-2.48, $P$=0.013) effects but no significant interaction between the two ($P$=0.40). These results recapitulate the linear mixed effects analysis in the main results, emphasizing both that participants tend to harvest longer in the short travel time blocks and that OUD participants do so more than controls on any given trial (i.e., regardless of travel time).

To test the marginal effect of lifetime opioid use on the probability of exiting the current patch at each trial, we repeated the above generalized linear mixed effects analysis in an OUD-only sub-analysis. Controlling for age, we found significant effects for travel time ($B$=-0.03, 95% $CI$ [-0.07, -0.002], $t_{18,311}$=-2.08, $P$=0.037) and lifetime opioid use ($B$=-0.02, 95% $CI$ [-0.05, -0.002], $t_{18,311}$=-2.16, $P$=0.03; two influential outliers excluded, Cook's $d$>0.07), but again no significant interaction effect ($P$=0.16), suggesting that increased lifetime opioid use is associated with reduced odds of exiting a patch regardless of travel time.

*Neuroimaging results: region-of-interest analysis using varying mask probability thresholds*

To examine the robustness of our region-of-interest definitions and analyses reported in the main text (see main text **Results** and **Fig. 2**), we repeated the analyses with participants' exit thresholds using probabilistic masks for the ventral tegmental area (VTA) and locus coeruleus (LC) defined at three different thresholds that have been previously reported in the literature: 0.05, 0.25, and 0.5 (most stringent). For each, we created binary masks using the SPM12 *imcalc* function on the probabilistic VTA and LC anatomical masks (1, 2). We then obtained each participant's mean neuromelanin signal contrast ratio by averaging over the voxels in the binary mask, excluding voxels with contrast ratios<0. These values for each participant were subsequently used in linear mixed effects analyses predicting signed deviation of participant's exit threshold from the MVT optimal threshold, as described in the main text. **Table S1** shows the results of these analyses including the regions of interest as predictors of participants' behavior, one for each mask probability threshold. We found that our results were robust to varying the mask probability threshold, with only increased neuromelanin signal in the VTA being significantly related to less overharvesting.

*Neuroimaging results: voxel-wise analysis in the LC*

Like for the VTA (see main text **Results** and **Fig. 3**), we examined the distribution of voxels within the entire LC that showed a positive relationship with foraging behavior. Unlike the region-of-interest analyses, this approach does not assume a homogeneous pattern across the LC, allowing for laterality or subregion effects to emerge as observed previously for LC neuromelanin in relation to other cognitive functions (3). Using a mask probability threshold of 0.05, 455 voxels were extracted for each participant ($n$=53). Voxels with a contrast ratio value less than the 1$^{st}$ and more than the 99$^{th}$ percentile across all participants were excluded. We then fit a robust regression to each voxel in the LC, predicting its signal contrast from behavior (mean signed deviation in participants' exit thresholds from the MVT optimal threshold), age, and repetition time of the scan acquisition, as described in the main text. Subsequently we obtained voxels that had a positive relationship to behavior ($\alpha<0.05$, uncorrected) and found that 15 voxels or less than 4% of the total met these criteria. These voxels were primarily located on the right side of the LC but did not demonstrate any meaningful clustering behavior.

*Neuroimaging results: association with decision noise and task engagement*

In accordance with prior animal work in which increased tonic norepinephrine correlated with performance decrements (4) and task disengagement leading to increased decision noise and earlier patch leaving in a similar patch-foraging task (5), we assessed for diagnostic group differences in these additional behavioral variables as well as the relationship to neuromelanin signal. For each participant, we computed as an index of decision noise the standard deviation on their trial-by-trial exit thresholds within each travel time block (short, long), while the number of late-response warnings (times when a participant failed to indicate a harvest-or-exit decision within the allotted time) were used as indices of task disengagement. Although our task was designed such that faster or slower response times could not affect earnings or the optimal choice policy so long as a response was registered in time (see main text **Methods**), late-responses could because they were effectively experienced as unforced timeouts.

There were no significant diagnosis, travel time, or diagnosis by travel time interaction effects on decision noise ($t_{75}<|0.39|$, $P>0.69$; linear mixed effects analysis predicting the standard deviation on participants' exit thresholds from diagnosis and travel time as fixed effects, with random slopes and intercepts by participant). Similarly, the frequency of late-response warnings did not differ between controls and participants with OUD ($t_{73}=-0.57$, $P=0.57$), with participants failing to indicate a decision within the allotted time in less than 5% of all decision opportunities. While response times tended to be faster overall in controls (295±86 ms) than in OUD participants (333±92 ms), this difference also did not reach significance ($t_{73}=-1.85$, $P=0.07$). These data indicate that lower-level factors such as task/attentional engagement do not explain the overharvesting bias observed in addiction. We also did not find evidence that neuromelanin signal in either the LC or VTA was related to decision noise or the frequency of late-responses (all $R<|0.17|$, $P>0.23$, controlling for age and repetition time of the scan acquisition), further suggesting that the observed relationship between increased VTA neuromelanin signal and less overharvesting also is not mediated by an effect on these lower-level factors.

### Supplemental Figures and Tables

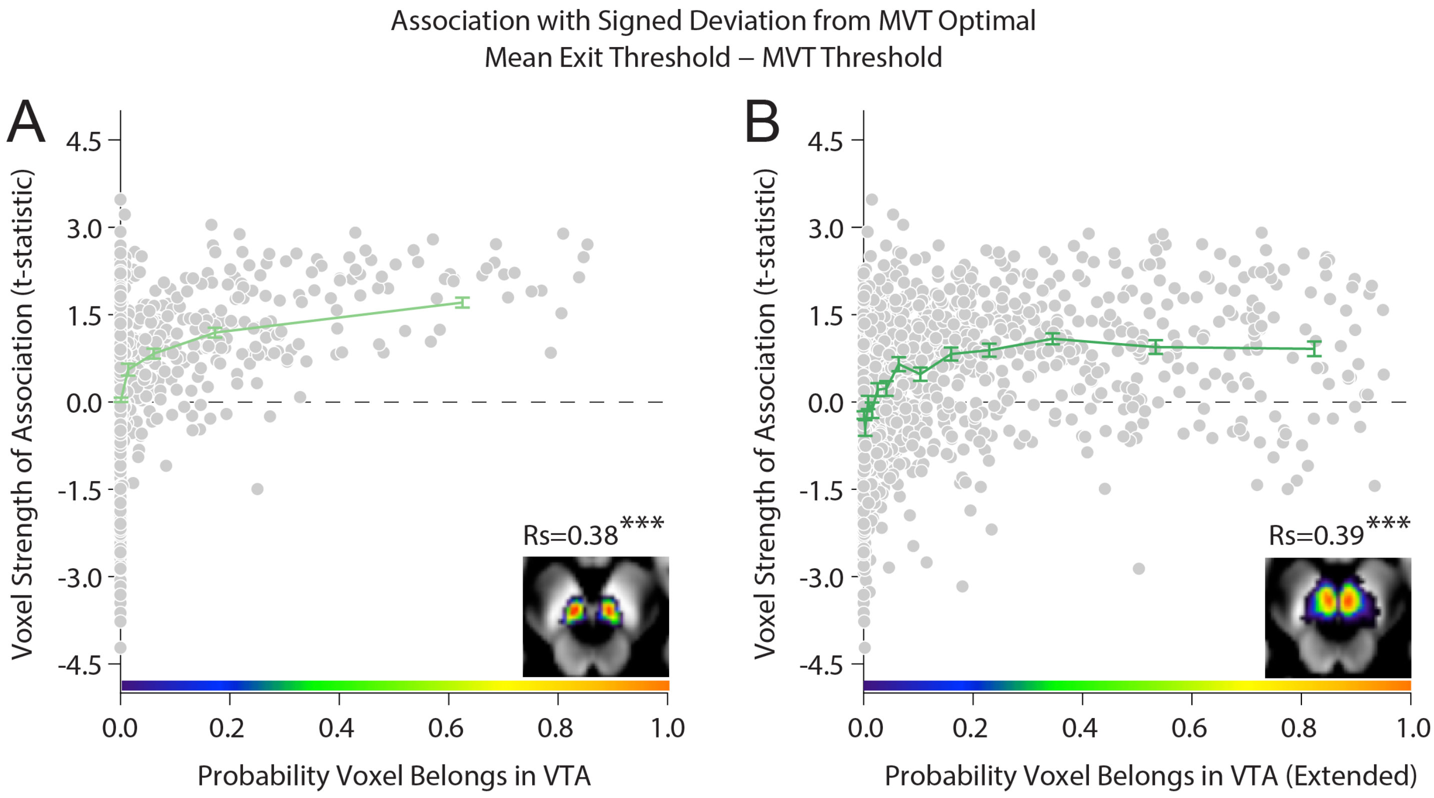


**Fig. S1. Relationship between a voxel's probability of belonging in the ventral tegmental area and its strength of association with foraging behavior (t-statistic).** Scatter plot of all voxels in the (**A**) probabilistic ventral tegmental area (VTA) mask used in the primary analyses (1) and (**B**) probabilistic mask using an alternative anatomical definition of the VTA that extends to the parabrachial nucleus (6), showing that voxels more strongly related to less overharvesting had higher probability of belonging to the VTA (controlling for age and repetition time [TR] of the scan acquisition). Inset shows the VTA masks at MNI coordinate *z*=-16 overlaid on the group average neuromelanin contrast ratio map (see main text). Green lines show quantiles (evenly spaced probability bins) for visualization purposes. The reported Spearman-*R* values use un-binned data. MVT, marginal value theorem; VTA, ventral tegmental area. *** $P<1.0\times10^{-307}$.

**Table S1. Association of neuromelanin signal in the ventral tegmental area and locus coeruleus with signed deviation of exit thresholds from MVT optimal, at different region-of-interest probabilistic mask thresholds***

| Region | Mask Probability Threshold | ***B***† | ***SE*** | ***t*–stat** | ***df*** ‡ | ***P*–value** | **95% *CI*** |
|---|---|---|---|---|---|---|---|
| VTA | 0.05 | 0.35 | 0.15 | 2.24 | 52.87 | 0.029 | [0.04, 0.66] |
| | 0.25 | 0.31 | 0.11 | 2.79 | 52.99 | 0.007 | [0.09, 0.54] |
| | 0.50 | 0.24 | 0.09 | 2.77 | 53.07 | 0.008 | [0.07, 0.41] |
| LC | 0.05 | 0.07 | 0.18 | 0.38 | 52.88 | 0.70 | [-0.29, 0.42] |
| | 0.25 | 0.09 | 0.16 | 0.60 | 52.85 | 0.55 | [-0.22, 0.40] |
| | 0.50 | 0.08 | 0.14 | 0.55 | 52.89 | 0.59 | [-0.20, 0.36] |

* Results of linear mixed-effects regressions including random intercepts and random slopes for travel time by participant and age, repetition time [TR] of the scan acquisition, and mean neuromelanin signal in each region of interest as fixed effects;
† Unstandardized coefficient;
‡ Degrees of freedom computed using Satterthwaite approximation.

**Table S2. Association of neuromelanin signal in dopaminergic subregions and the noradrenergic locus coeruleus with signed deviation of exit thresholds from MVT optimal ***

| Region | ***B***[†] | ***SE*** | ***t*–stat** | ***df*** [‡] | ***P*–value** | **95% *CI*** |
|---|---|---|---|---|---|---|
| VTA | 0.48 | 0.16 | 3.02 | 53.45 | 0.004 | [0.16, 0.79] |
| SNc | -0.36 | 0.38 | -0.96 | 53.39 | 0.34 | [-1.11, 0.39] |
| SNr | 0.35 | 0.33 | 1.07 | 53.29 | 0.29 | [-0.31, 1.02] |
| LC | -0.42 | 0.24 | -1.76 | 52.45 | 0.08 | [-0.91, 0.06] |

[*] Results of a linear mixed-effects regression including random intercepts and random slopes for travel time by participant and age, repetition time [TR] of the scan acquisition, and mean neuromelanin signal in the listed regions of interest as fixed effects;
[†] Unstandardized coefficient;
[‡] Degrees of freedom computed using Satterthwaite approximation.

**Table S3. Association of neuromelanin signal in dopaminergic subregions and the noradrenergic locus coeruleus with lifetime opioid use ***

| Region | *B*[†] | *SE* | *t*–stat | *df* | *P*–value |
|---|---|---|---|---|---|
| VTA | -2.85 | 1.31 | -2.17 | 18 | 0.04 |
| SNc | 3.91 | 3.08 | 1.27 | 18 | 0.22 |
| SNr | -1.17 | 2.62 | -0.45 | 18 | 0.66 |
| LC | 1.52 | 1.90 | 0.80 | 18 | 0.44 |

[*] Results of a linear regression predicting lifetime years of opioid use within opioid use disorder participants from age, repetition time [TR] of the scan acquisition, and mean neuromelanin signal in the listed regions of interest;
[†] Unstandardized coefficient.

**Table S4. Association of neuromelanin signal in dopaminergic subregions and the noradrenergic locus coeruleus with opioid use disorder class ***

| Region | *B*[†] | *SE* | *t*–stat | *df* | *P*–value |
|---|---|---|---|---|---|
| VTA | -0.53 | 0.26 | -2.08 | 46 | 0.04 |
| SNc | 0.67 | 0.54 | 1.24 | 46 | 0.22 |
| SNr | -0.41 | 0.45 | -0.91 | 46 | 0.36 |
| LC | 0.65 | 0.34 | 1.92 | 46 | 0.06 |

[*] Results of a logistic regression predicting opioid use disorder diagnosis class (1 or 0) from age, repetition time [TR] of the scan acquisition, and mean neuromelanin signal in the listed regions of interest;
[†] Unstandardized coefficient.
